# Applying an interpretable machine learning approach to assess intraspecific trait variation under landscape-scale population differentiation

**DOI:** 10.1101/2023.04.07.536012

**Authors:** Sambadi Majumder, Chase M. Mason

**Affiliations:** Department of Biology, University of Central Florida, Orlando, FL

**Keywords:** Boruta, ecophysiology, feature selection, gradient boosting machine, *Helianthus*, multidimensional, random forest, accumulated local effects

## Abstract

**Premise:** Here we demonstrate the application of interpretable machine learning methods to investigate intraspecific functional trait divergence using diverse genotypes of the wide-ranging sunflower *Helianthus annuus* occupying populations across contrasting ecoregions - the Great Plains versus the North American Deserts.

**Methods:** Recursive feature elimination was applied to functional trait data from the HeliantHome database, followed by the application of Boruta to detect traits most predictive of ecoregion. Random Forest and Gradient Boosting Machine classifiers were then trained and validated, with results visualized using accumulated local effects plots.

**Key Results:** The most ecoregion-predictive functional traits span categories of leaf economics, plant architecture, reproductive phenology, and floral and seed morphology. Relative to the Great Plains, genotypes from the North American Deserts exhibit shorter stature, fewer leaves, higher leaf nitrogen, and longer average length of phyllaries.

**Conclusions:** This approach readily identifies traits predictive of ecoregion origin, and thus functional traits most likely to be responsible for contrasting ecological strategies across the landscape. This type of approach can be used to parse large plant trait datasets in a wide range of contexts, including explicitly testing the applicability of interspecific paradigms at intraspecific scales.

## INTRODUCTION

Plants exhibit a wide range of variation in traits that shape their resource allocation and environmental interactions, influencing their overall fitness through growth, survival, and reproduction (Caruso et al., 2020). These “functional traits” are morphological, physiological, phenological, or chemical characteristics that vary not only among species under macroevolutionary divergence, but also among populations within species in response to variation in environmental conditions across ranges (Violle et al., 2007; Díaz et al., 2013; Caruso et al., 2020). Interspecific variation in functional traits has been used by researchers to describe plant ecological strategies, resulting in trait-based paradigms like the competitive-stress tolerant-ruderal framework (CSR; Grime, 1977), the leaf-height-seed scheme (LHS; Westoby at al., 1998), and the leaf economics spectrum (LES, Wright et al., 2004), later expanded to stems, roots, flowers, and whole plants (Baraloto et al., 2010; Mommer et al., 2012; Reich, 2014; Roddy et al., 2020). While these trait-based frameworks for ecological divergence have largely focused on among-species variation, intraspecific variation (both within and among plant populations) has been demonstrated to account for between one-quarter and one-third of all trait variation within and among communities (Albert et al., 2010; Siefert et al., 2015) and to strongly impact plant growth responses to environmental factors (Violle et al., 2012; Laforest-Lapointe et al., 2014; Brouillette et al., 2014; May et al., 2017; Araújo et al., 2021) as well as total biodiversity (Raffard et al., 2019). For example, the LHS paradigm applied to Scots pine (*Pinus sylvestris*) across its range in northern Spain identified that traits like maximum height and leaf nitrogen were critical drivers of tree growth rate across climate gradients (Laforest-Lapointe et al., 2014). The LES paradigm has similarly been applied at the intraspecific scale within many different species, describing local adaptation across geographic ranges in wild plants like Arabidopsis (Vasseur et al., 2012); western sunflower (*Helianthus anomalus*; Brouillette et al., 2014), and holly oak (*Quercus ilex*; Niinemets, 2015), as well as in crops like soybean, rice, wheat, maize, and even coffee (Martin et al., 2017; Martin et al., 2018; Xiong et al., 2018; Hayes et al., 2019). In most cases, trait-based ecological paradigms use small sets of traits to capture major aspects of plant multivariate trait variation that have perceived *a priori* ecological importance, mainly derived from existing knowledge of plant physiology across assorted study systems. This trend in part derives from a community goal of developing an approximate consensus list of plant traits reflecting ecological strategies ranked by their ‘importance’ (Westoby et al., 2002), which facilitates meta-analysis and synthesis across studies as well as the forecasting of future vegetation dynamics under changing climate conditions (Westoby, 1998). However, there is no guarantee that such ‘consensus’ lists of traits reflect the most evolutionarily important or divergent traits under local adaptation among populations of a species, given the multivariate nature of selective pressures arising from the abiotic or biotic environment. Put simply, approaches based on few-trait paradigms like the CSR, LHS, or LES may overlook critical functional traits with high ecological importance, and this is especially likely at the intraspecific scale where trait variation is usually only a small portion of global or community-level interspecific variation (Klimešová et al., 2015; Niinemets, 2015; Anderegg et al., 2018).

Conversely, the collection of large trait datasets (dozens to hundreds of traits) at the intraspecific level offers a major opportunity to explicitly test the generality of these few-trait paradigms and may be a route to validating their utility across scales or circumscribing when these paradigms are most and least useful (e.g., by phylogenetic lineage, growth form, etc). Furthermore, such approaches have the ability to identify important but uncommonly assessed plant traits that may have been overlooked by researchers due to inconvenience of measurement, biases of human perception, lineage-related predispositions of focus, or historical happenstance.

In an era of large-scale access to both datasets and computational resources, trait databases are becoming a vital component of advancing plant science and biodiversity research. Widely known databases include tabular trait datasets (e.g., TRY; Kattge et al, 2020), as well as occurrence data and digitized herbarium specimen images (e.g., the Global Biodiversity Information Facility; GBIF 2021b). In this work, we utilize functional trait data from the HeliantHome database (Bercovich et al., 2022) derived from common garden phenotyping of hundreds of genotypes of wild common sunflower (*Helianthus annuus* L.) belonging to populations across the native range of this widespread species. Common garden experiments are a classic approach to understanding local adaptation, assessing the extent of genetic differentiation in phenotypes among a set of genotypes spanning populations (e.g., Clausen et al., 1940). From a quantitative genetics perspective, comparing plant genotypes under near-identical environmental conditions strongly reduces environmentally-driven plastic variation (E) in plant traits relative to in situ field phenotyping across ranges and permits the estimation of the genetic components (G and GxE) of trait variation – as well as a range of other useful inferences (De Villemereuil et al., 2016; Schwinning et al., 2022). Here we specifically utilize phenotypic data associated with genotypes derived from source populations spanning the two largest Level I ecoregions this species occupies – the Great Plains and the North American Deserts (Figure 1; U.S. Environmental Protection Agency 2010). The progenitor of cultivated sunflower, the wild common sunflower occupies a substantial native range of least four million square kilometers and is known to exhibit broad diversity in phenotypes across this geographic expanse (Todesco et al., 2019; Todesco et al., 2022; Bercovich et al., 2022). This intraspecific variation likely reflects local adaptation of functional traits to facilitate growth and survival across gradients of precipitation, soil fertility, and other environmental factors (Heiser et al., 1969). Wild common sunflower is therefore a useful system to demonstrate the application of interpretable descriptive and predictive machine learning (ML) models using structured tabular trait data to make inferences about geographic differentiation in plant traits. Specifically, we here attempt to identify a core set of traits that are most divergent between populations native to contrasting biomes, here the Great Plains versus the North American Deserts. Our approach allows for the ranking of plant traits based on their relative “importance” in classifying genotypes to the correct source ecoregion, highlighting dimensions of high intraspecific divergence in multivariate trait space. It also allows us to ascertain how plant phenotypes of populations growing in the Great Plains differ from those found in the North American Deserts. This provides insight into intraspecific trait diversification along key functional trait axes and facilitates within-species comparison of ecological strategies.

**Figure 1.**
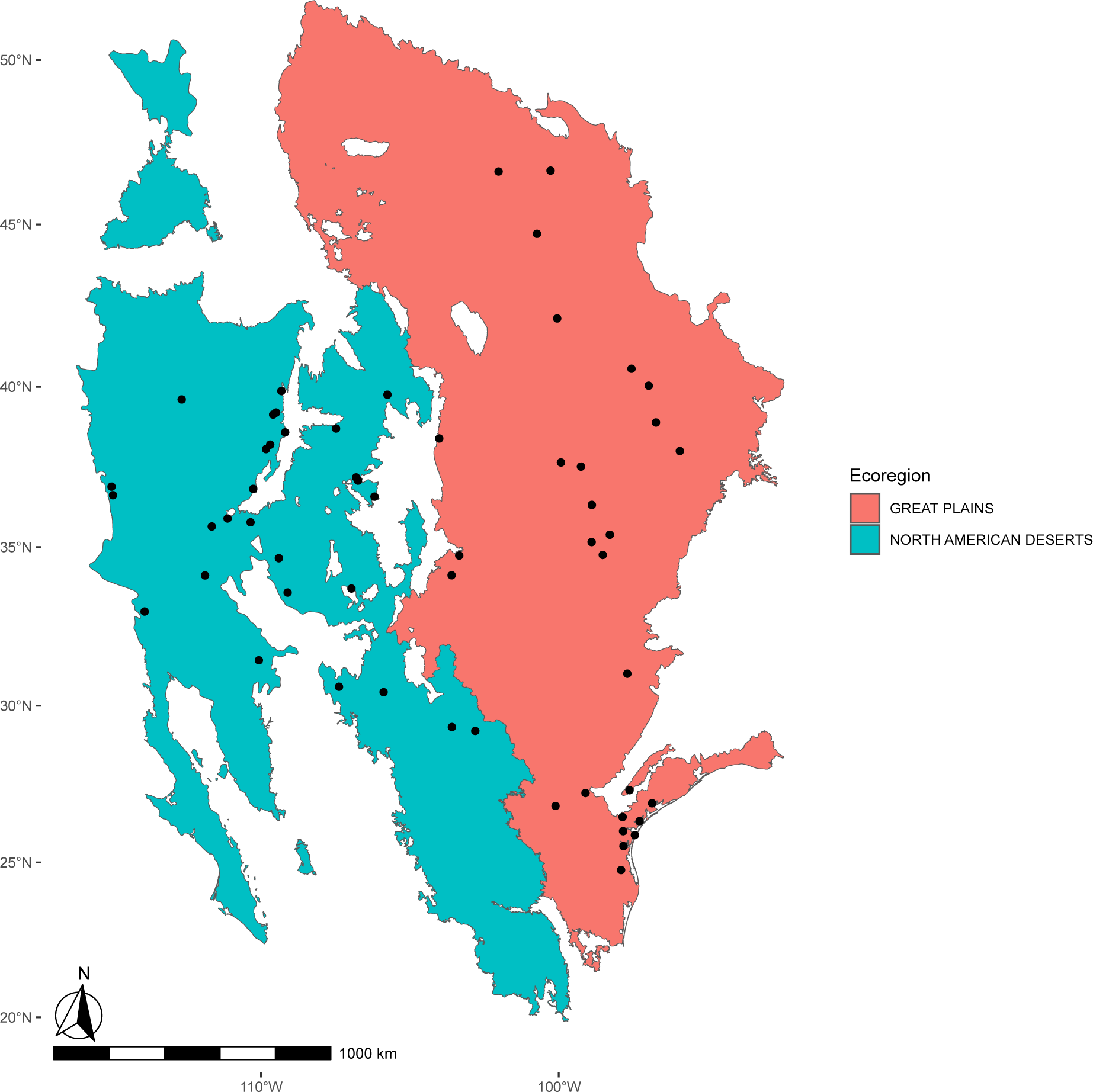
Map of the two Level I Ecoregions used in this study (the Great Plains, in pink, and The North American Deserts, in blue) with points representing the locations of the populations of *Helianthus annuus* used in this study, as derived from the HeliantHome database.

## METHODS

### Software and programming language

All data compiling, cleaning, exploration, visualizations, and modeling workflows were formulated using packages in the R programming environment (R Core Team 2022). A laptop containing 8GB RAM. All code can be accessed through the project GitHub (https://github.com/SamMajumder/Applying-XAI-approaches-to-ecology/tree/main).

### Study system, study location, data source and data preparation

From all available genotypes of *Helianthus annuus* within the public HeliantHome database (Bercovich et al., 2022), those occurring within the Level I ecoregions of the Great Plains and North American Deserts were extracted by cross-referencing the geographic coordinates of each source population with a shapefile of Level I ecoregions sourced from the United States Environmental Protection Agency (U.S. Environmental Protection Agency, 2010). These two ecoregions were selected because they contain the majority of *H. annuus* populations, with other ecoregions having too few populations to have sufficient statistical power for analysis. For these genotypes, all available functional trait data was also obtained from HeliantHome. This data contains a wide range of traits reflecting plant architecture, reproductive phenology, tissue chemistry, and morphology of leaves, stems, inflorescences, and seeds, with trait values associated to one or more genotypes within each population. A combined dataset was generated containing the population coordinates, the categorical ecoregion assignment, the genotypes within these populations, and the corresponding functional trait data for each genotype by leveraging R packages such as *sf* (Pebesma, 2018), *jsonlite* (Ooms, 2014) and the *tidyverse* suite of packages (Wickham et al., 2019). This combined dataset contained 88 traits from 464 genotypes belonging to 49 populations (Figure 1), where each individual observation was a unique genotype. The dataset was then examined to identify the percentage of missing values for each trait (Figure S1). One of the traits, the peduncle length of the first flower, had 100% missing values and was removed from the dataset. All other traits had missing values at a rate of either 27% (seed traits), or <5% (all other traits). The combined dataset, now containing 87 traits (Table S1), was then randomly divided into a training dataset and a testing dataset. Seventy percent of the individuals in the data were used for training, with the remaining thirty percent used for testing to evaluate the predictive models on unseen data. Missing data values were imputed by using the proximity matrix from a Random Forest (RF) algorithm (Breiman, 2001) through the R package *randomForest* (Liaw and Wiener 2002). This imputation of missing values was performed on the training data (Table S2) and testing data (Table S3) separately to prevent data leakage (Figure 2).

**Figure 2.**
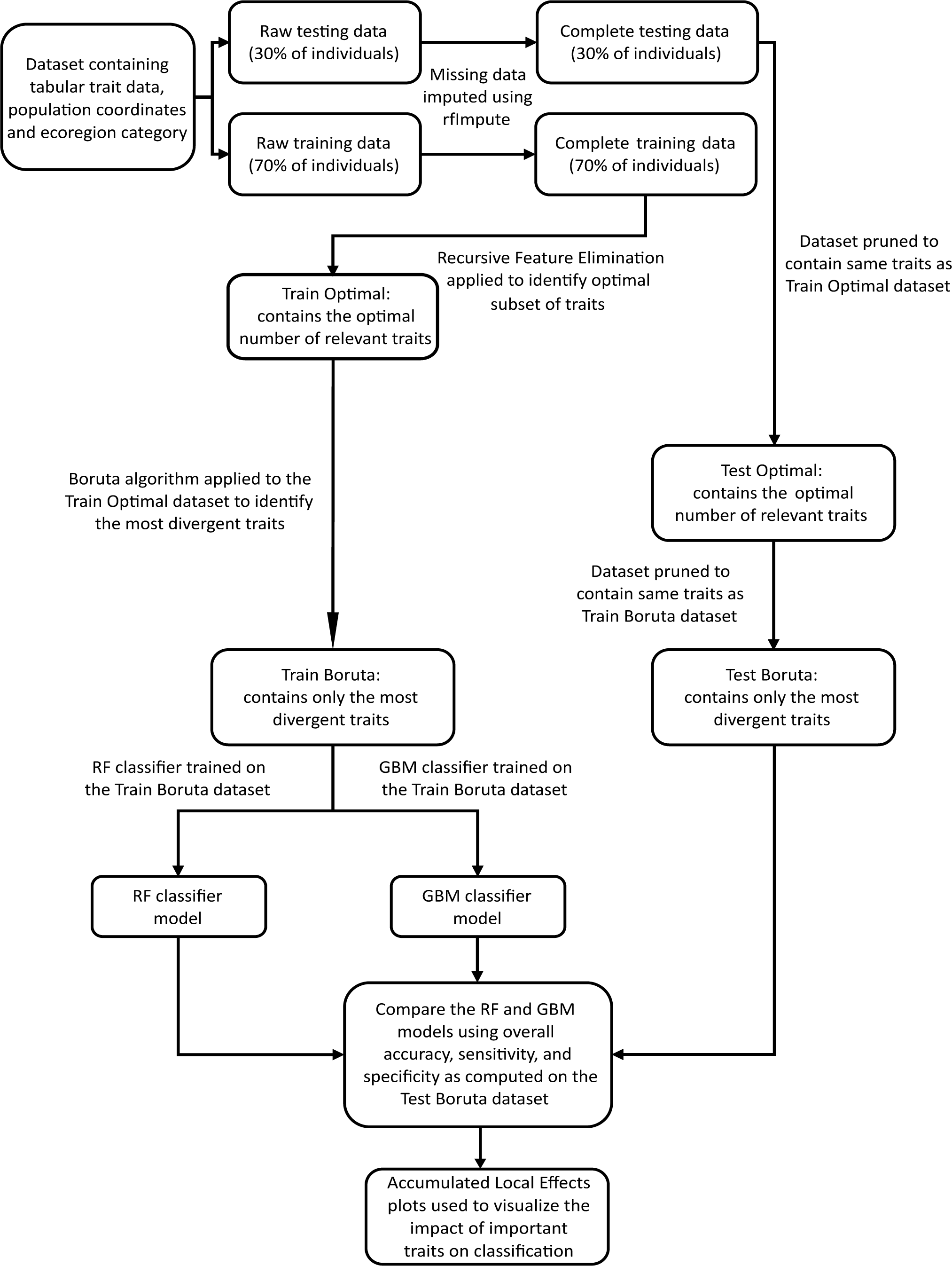
Complete workflow for the machine learning analysis implemented in this study. The input dataset contained HeliantHome database functional trait data from *Helianthus annuus* genotypes, and corresponding geographical coordinates from source populations across the Great Plains and The North American Deserts ecoregions. The data was divided into a training and a test dataset by random sampling, whereby 70% of the data was used for training and 30% was used for testing. Missing data in both the training and the test datasets were imputed separately, to avoid data leakage, using the random forest algorithm. This was implemented using the “rfImpute” function from the *radomForest* R package (Liaw and Wiener, 2002). Recursive feature elimination (RFE) was then applied on the training data to identify the optimal subset of relevant traits (i.e., those which are potentially good predictors of ecoregion origin). The training dataset was then reduced to contain only the optimal subset of traits (generating ‘Train Optimal’), and the same traits were also discarded from the test dataset (generating ‘Test Optimal’). Next, the Boruta algorithm was applied on the ‘Train optimal’ dataset to identify the most divergent traits (generating ‘Train Boruta’), and the same traits were retained in the parallel test dataset (‘Test Boruta’). To validate the findings of RFE and Boruta, two classifiers, Random Forest (RF) and Gradient Boosting Machine (GBM), were trained separately on the ‘Train Boruta’ dataset, and evaluated on the ‘Test Boruta’ dataset. This approach allowed us to determine how well the traits deemed both most important by RFE and most divergent by Boruta predict ecoregion origin, and by extension enabled us to understand whether these traits reflect phenotypic divergence between populations from these contrasting environments. Accumulated local effects (ALE) plots were then used to evaluate the specific impact on prediction probability of variation in trait values for each of the eight most important traits.

### Classification algorithms used for feature selection and subsequent predictive modeling

Throughout this work we utilize two tree-based ensemble ML classification algorithms, Random Forest (RF) and Gradient Boosting Machine (GBM), in various ways. We use variable importance values calculated by RF or RF-based frameworks to identify the most phenotypically divergent functional traits between two sets of populations from the ecoregions assessed in this study (i.e., feature selection). Additionally, we trained two predictive models to validate the variables determined to be ‘important’ during feature selection. RF creates several decision trees, and each tree is trained on a bootstrapped version of the original training data (Pal 2005; Valletta et al., 2017). The bootstrapped dataset is created by randomly sampling from the data with replacement. The predictions made from each decision trees are then averaged across all trees.

The “out of bag” (OOB) data, the portion left out when creating the bootstrapped dataset, is used to evaluate the predictive capabilities of the trained ensemble classifier (Cutler et al., 2012; Valletta et al., 2017). GBM also creates several decision trees during the training process, but differs from RF in that each tree in the ensemble seeks to reduce the error of the tree that came before it, sequentially improving the prediction accuracy of the overall model (Friedman, 2001).

### Identifying the optimal subset of strongly ecoregion-divergent traits and their impacts on classification

As a backward elimination process, Recursive Feature Elimination (RFE) selects important variables within a dataset by recursively removing unimportant variables through a process of multiple models built on the training data using a ML classifier chosen by the user (Guyon et al., 2008). In this study, we used RF (Breiman, 2001) as our classifier within RFE. Variables are ranked based on their overall importance to the prediction specified (here binary classification) and with each iteration of model building, the lowest ranked variable is removed from the dataset. Importance of variables were calculated using Mean Decrease of Accuracy (MDA) (Breiman, 2001). To accomplish this, first a baseline prediction accuracy is calculated on the OOB data. Then the value of a variable is permuted, which causes a difference in the prediction accuracy and this difference is recorded. This accuracy difference is averaged and normalized across all trees in the ensemble (Kuhn, 2008). Conceptually, according to MDA, the importance of a variable is the decline of prediction accuracy seen after permutation. This process is repeated for all variables in the training dataset in order to compute the variable importance value for each. Here, RFE was used to identify the functional traits that are most important for reliable classification of genotypes to plains versus deserts. Put another way, RFE was used to identify the optimal subset of traits most consistently exhibiting clear intraspecific divergence between the two ecoregions and was implemented in this study using the R package called caret (Kuhn, 2008). Only the traits identified as “important” by RFE were retained in the training dataset, and the rest were discarded (Figure 2).

To identify the most strongly divergent traits among the list of optimal traits in relation to phenotypic divergence, the Boruta algorithm (Kursa and Rudnicki, 2010) was applied on the training dataset containing only the variables deemed as “important” by RFE (Figure 2). As per the *all-relevant* problem (Nilsson et al., 2007; Kursa and Rudnicki, 2010) some variables (or features) in a dataset are more influential than others within a particular classification task. To demystify the black-box nature of ML-aided classification tasks, a good understanding of the varied influences of features in a given dataset on the overall classification outcome is necessary (Kursa and Rudnicki, 2010). The Boruta algorithm, an RF-based algorithm, was implemented here by using the *Boruta* R package (Kursa and Rudnicki, 2010). Boruta is useful to determine the most strongly relevant, weakly relevant, and redundant variables in a dataset by using a permutation-based method to compute variable importance, one that differs from MDA. While MDA permutes the values of the variables in the OOB data, Boruta first duplicates the entire training dataset where the values of the duplicated variables are obtained by randomly permuting the original values of the variables of the training dataset. Thus, the dataset now contains the variables from the original training dataset, referred to as “original variables” and their duplicated counterparts are referred to as “shadow variables”. The RF algorithm is then applied to this combined dataset and the importance scores of both the “original variables” and “shadow variables” were calculated. If the importance score of a given variable from the original training dataset (“original variables”) is more than the maximum importance score among the shadow variable then it is denoted as strongly relevant, if that is not the case then the variable is weakly relevant. After the Boruta algorithm was applied, the strongly relevant traits in relation to ecological divergence were retained in the training dataset while the rest were removed. This reduced version of the training dataset was used to train a Random Forest (RF) and a Gradient Boosting Machine (GBM) classifier (Figure 2). The same traits were also retained in the test data as subsequent model validation requires a dataset containing the same exact variables on which the classifier was trained.

### Training machine learning classifiers

The two predictive models created with RF and GBM were binary classifiers, where the two classes were the ecoregions of Great Plains (“0”) and North American Deserts (“1”). The predictive capabilities of the two models to successfully classify each class (ecoregion) were compared using their respective sensitivity and specificity scores. In this implementation, sensitivity indicates the ability of the model to correctly classify genotypes from the Great Plains (true “0”), whereas specificity indicates the ability of the model to correctly classify genotypes from the North American Deserts (true “1”). These metrics were calculated after the models were applied to the test dataset and were thus also used to ascertain whether the models were overfitting on the training data, thus evaluating the validity of the traits identified as ‘important’. This reduced the likelihood of drawing erroneous biological conclusions due to overfitting and provides confidence in the traits identified by Boruta to be relevant to phenotypic divergence between ecoregions. Accumulated local effects (ALE) plots (Apley et al., 2020) were used to describe the impact of trait values on the overall prediction of the ML classifier (RF), as ALE plots are robust to datasets where features are correlated with each other (Molnar, 2021) as is typical of plant functional trait data.

## RESULTS

A total of 51 traits were identified by RFE as important to the accurate classification of source ecoregion, and therefore are divergent between the two sets of populations from the Great Plains and North American Deserts (Table S4; Figure 3). This list included leaf morphological traits like shape descriptors, and leaf physiological traits related to the leaf economic spectrum, including leaf nitrogen content, leaf C:N ratio, specific leaf area (SLA), as well as total leaf RGB and the proportion of red and blue in leaf color – indicative of pigmentation including chlorophyll and anthocyanin content. Floral morphological traits were especially common, including flower head diameter, the size and shape of phyllaries, size and number of ligules (petals), and various morphometric ratios of floral parts. Whole plant architectural and phenological traits were also identified as important, such as number of primary branches, total leaf number, distance between the ground and first node (indicative of plant height at first branching), days to budding and flowering, and plant height and stem diameter at flowering.

**Figure 3.**
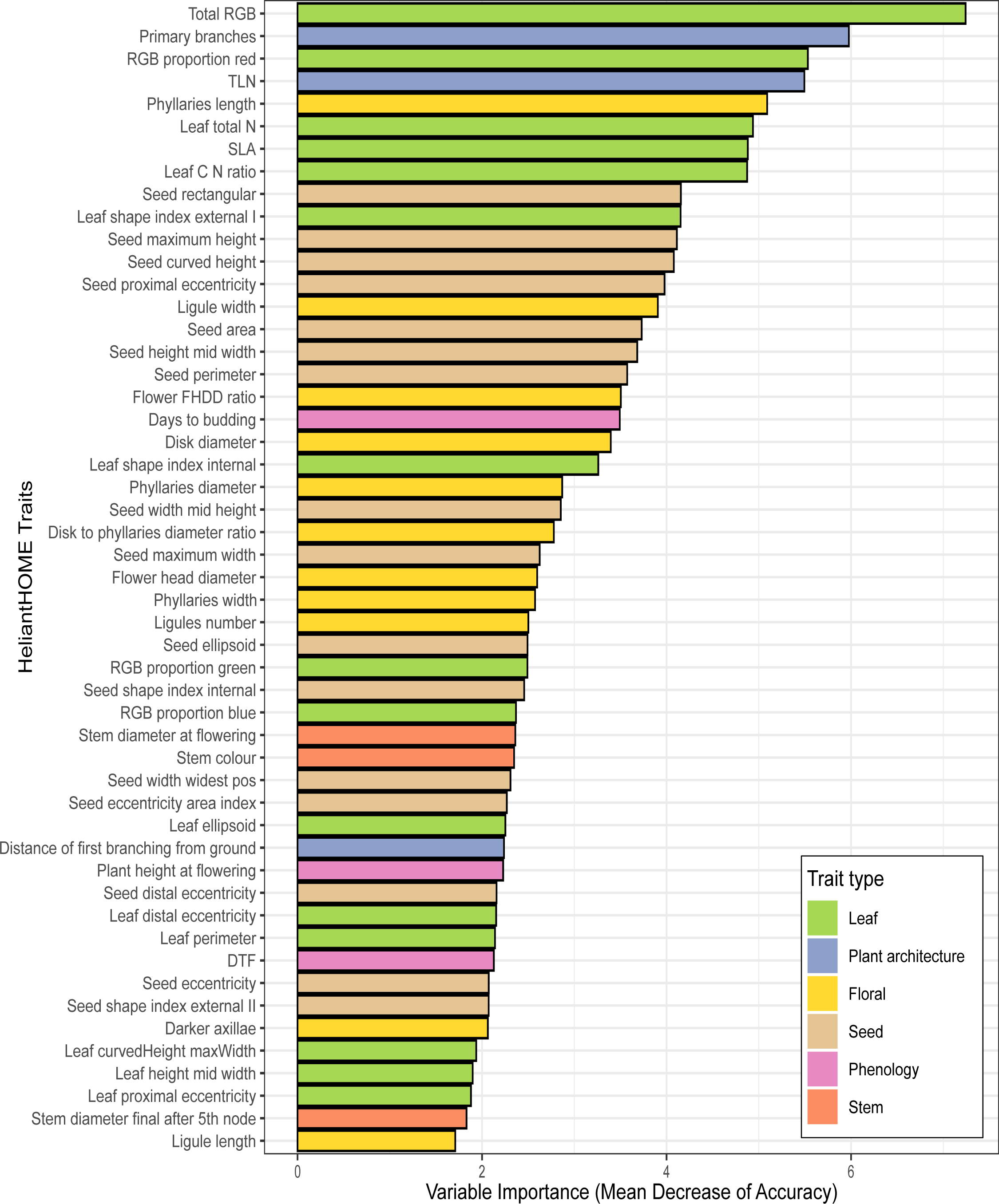
The 51 traits identified as part of the optimal subset of relevant traits, by recursive feature elimination, ranked by the order of their relative importance computed using Mean Decrease of Accuracy (MDA). A high MDA value indicates that intraspecific phenotypic divergence is stronger in relation to ecoregion, and these 51 traits are inferred to be traits at least minimally important to ecoregion divergence as compared to the other 36 traits in the full dataset which were not identified to be in the optimal subset.

Other important traits included a range of seed size and shape metrics, and pigmentation of stems and axillary buds. Conversely, 36 traits were not selected by RFE, indicating these traits are not divergent between ecoregions (Table S5). Among these traits are a range of leaf traits relating to overall leaf size and shape, carbon content, trichome density, and petiole color, floral morphology traits including peduncle length, certain seed morphology traits, and whole-plant traits like internode length and leaf initiation rate.

Subsequently, Boruta identified 35 traits as potentially strongly divergent (Table S6) between the two sets of populations (i.e., strongly delineating them in a multidimensional trait space). These traits were ranked in the order of importance, and the top eight traits in this list are total leaf RGB, number of primary branches, RGB proportion red, total leaf number, length of individual phyllaries, total leaf nitrogen, specific leaf area, and leaf carbon to nitrogen ratio. It is interesting to note that the ranking of traits was predominantly the same between the RFE and Boruta approaches with few exceptions.

Comparing the two classifiers, RF outperformed GBM when correctly classifying the genotypes from the North American Desert (specificity scores on the test dataset were RF: 0.88, GBM: 0.80), whereas the GBM outperformed RF in correctly classifying genotypes from the Great Plains (sensitivity scores on the test dataset were RF: 0.67, GBM: 0.78). Geographic distribution maps were used to visualize the number of genotypes per population that were correctly predicted by the two classifiers, one for GBM (Figure 4) and one for RF (Figure S2). RF incorrectly predicted all genotypes within populations ANN_44 (genotypes: ANN1231, ANN1232, ANN1233 and ANN1234, Ecoregion: The Great Plains), ANN_19 (genotypes: ANN0983 and ANN0986, Ecoregion: The Great Plains) and ANN_09 (genotype: ANN0886, Ecoregion: North American Deserts) (Table S7), while GBM performed a bit better, correctly predicting 1 out of 4 genotypes for ANN_44 (correct prediction of genotype: ANN1233), 1 out of 2 for ANN_19 (correct prediction of genotype: ANN0983), and 1 out of 1 for ANN_09 (Table S8). Interestingly, most populations with a high rate of misclassified genotypes appear to be located near ecoregion boundaries (Figure 4). As GBM showed reasonably high sensitivity and specificity values (able to classify both ecoregions reasonably well compared to RF) and was also able to correctly classify at least one genotype from each population, its prediction probabilities were chosen to compute ALE values and visualized in plots for several of the more important traits (Figure 5). Trait thresholds for phenotypic divergence between ecoregions can be readily identified – for example, leaf nitrogen has a threshold value of approximately 5.75%, leaf carbon-to-nitrogen ratio has a threshold value of around 11, and total leaf RGB has a threshold value of approximately 160 (Figure 5). These thresholds indicate higher concentrations of leaf nutrients and photosynthetic pigments within populations from the North American Deserts compared to populations from the Great Plains. With respect to whole plant architecture, populations from deserts seem to exhibit fewer leaves and primary branches at maturity when compared to populations from the plains, with approximate threshold values of 20 for both traits (Figure 5). Higher leaf construction cost, as indicated by low specific leaf area, is predictive of populations from the Great Plains, especially values below a threshold of 20 mm mg^−1^ (Figure 5). Populations from deserts also seem to have a higher proportion of the color red in their leaves, indicating ultraviolet-protective anthocyanin pigments, as well as longer phyllaries subtending the composite head (Figure 5). When visualizing the original distribution of univariate trait values for the top eight traits via raincloud plots (Figure 6), the directionality of trait differences as visualized with ALE plots is retained, but it becomes apparent that individual traits alone are insufficient to delineate ecoregions given the high phenotypic variation found within each. Only using multivariate approaches like those employed here is it possible to predict ecoregion origin with a high success rate and identify traits that have most consistently diverged between ecoregions such that they are important to the success of ecoregion classification.

**Figure 4.**
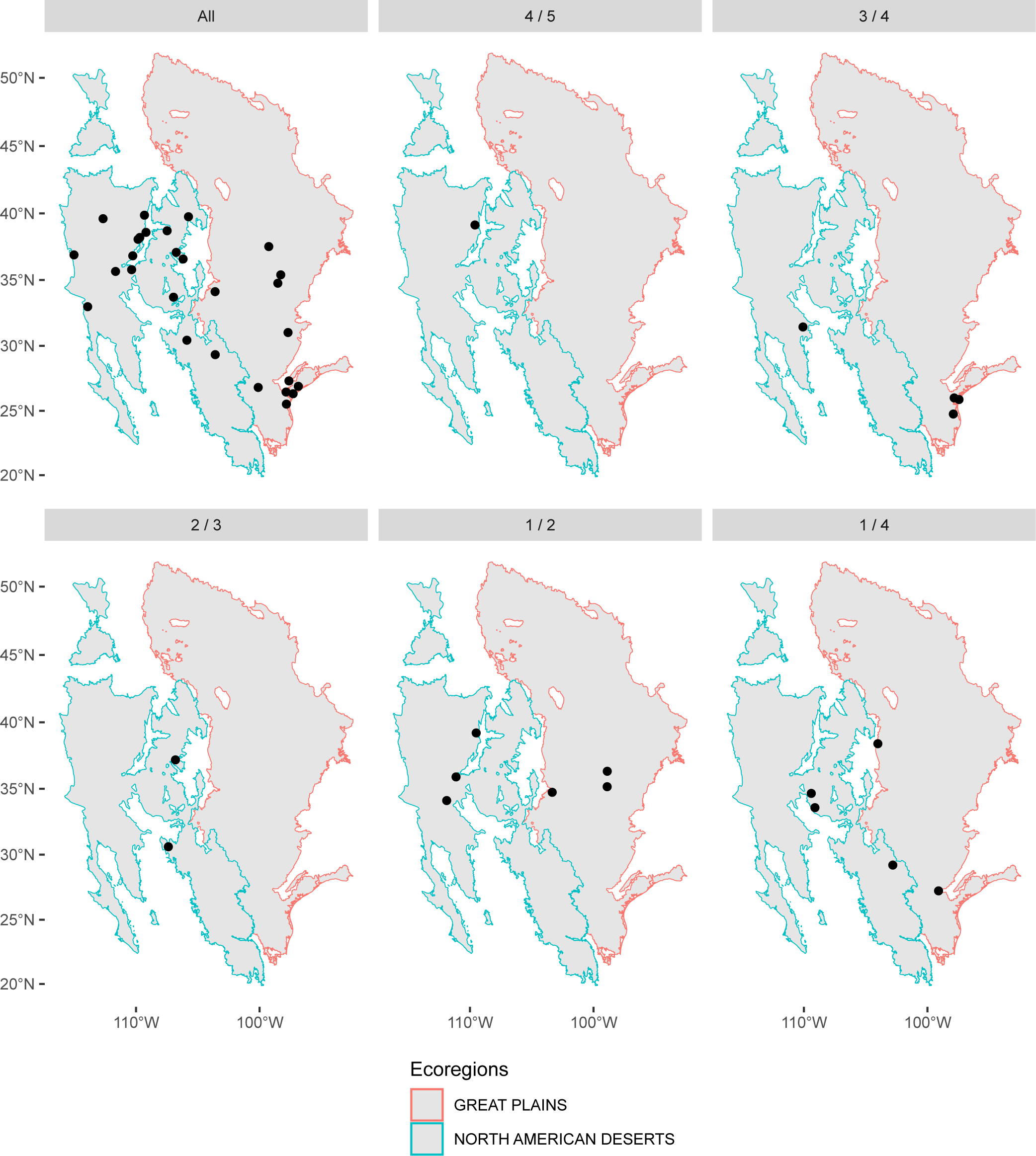
Visualizations of the predictions made by the Gradient Boosting Machine (GBM) classifier, where points represent populations of *H. annuus*. Each facet represents populations where a given proportion of genotypes was correctly predicted, arranged in descending order – all genotypes correctly predicted, four out of five, three out of four, two out of three, one out of two, and one out of four. In no populations were zero genotypes correctly predicted by GBM.

**Figure 5.**
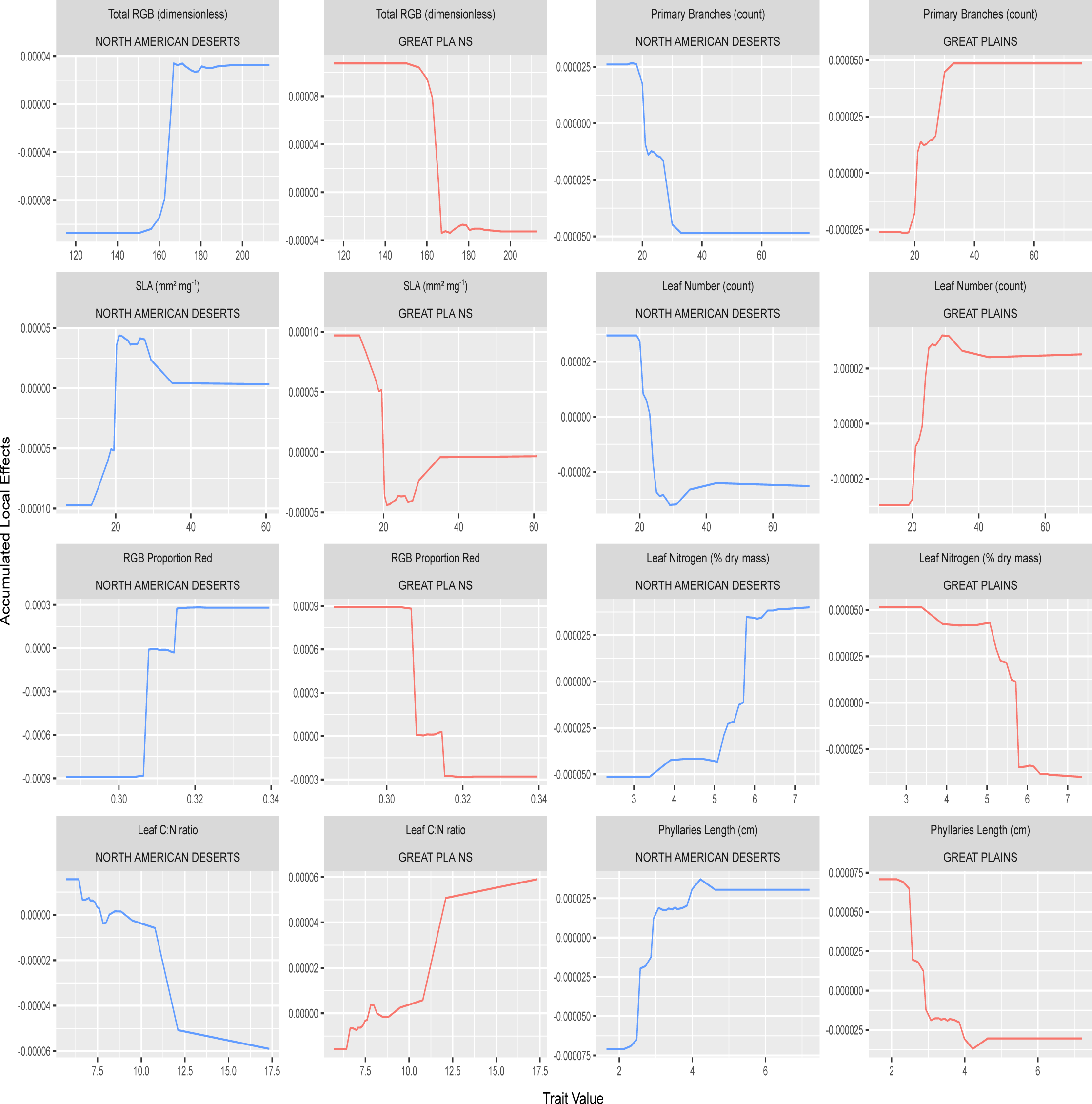
Accumulated local effects (ALE) calculated from the prediction probabilities generated by the GBM classifier and plotted for the eight traits with highest importance to ecoregion prediction within the optimal subset of relevant traits, as per RFE (Figure 3), and also part of the set of strongly ecoregion-divergent traits as per Boruta (Table S6). Top panels reflect ALE for The North American Deserts, while bottom panels reflect ALE values for the Great Plains. Individual plots show the impact of the trait value for a given trait on the prediction probability for a given ecoregion, with a positive ALE value favoring classification to the ecoregion, and a negative ALE value disfavoring classification to the ecoregion. ALE values equal to zero indicate threshold trait values between ecoregions.

**Figure 6.**
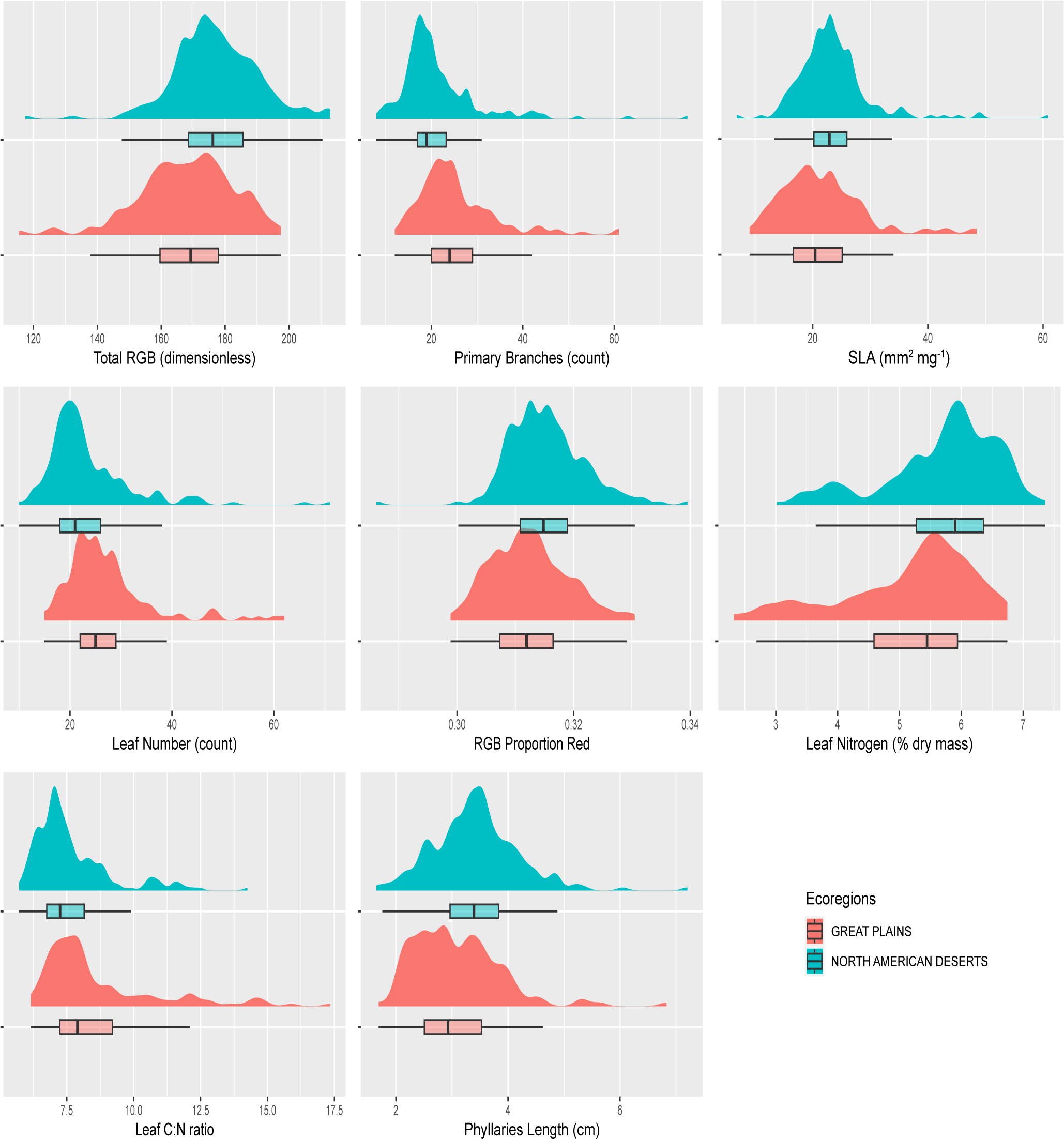
Univariate trait distributions for the eight traits with highest importance to ecoregion prediction. Distributions are smoothed histograms representing relative data density, while boxplots reflect the median and quartiles, with lines indicating 1.5 times the interquartile range on either side of the first and third quartile.

## DISCUSSION

Many studies have acknowledged the importance of intraspecific functional trait variability in evolutionary as well as ecological processes (Newton et al., 1999; Moran et al., 2016; Westerband et al., 2021) and attempts have been made to analyze this variation to more accurately understand functional diversity and complex community-based ecological interactions (Violle et al., 2012; Kuppler et al., 2016). Our ML-based approach to studying intraspecific divergence in functional traits allows us to implement an objective trait-first approach to examining intraspecific phenotypic divergence. We then attempt to analyze our findings through the lens of existing ecological paradigms and determine whether the major dimensions of phenotypic divergence fall within these existing trait-based frameworks. Our results indicate that the largest dimensions of intraspecific multivariate phenotypic divergence between ecoregions in *H. annuu*s involve leaf physiology, plant architecture, reproductive phenology, and certain floral, stem, seed, and leaf morphology traits (Table S6). Together, model results indicate that the LES paradigm (leaf investment and return) is a key dimension of divergence, given the high importance of construction costs, nutrient investment, and pigmentation, all drivers of mass-based photosynthetic rate and leaf-level net primary productivity (Wright et al., 2004; Mason et al., 2015). This axis of variation has been identified as important for intraspecific population divergence across environmental gradients within another annual sunflower species (Brouillette et al., 2014), as well as at the interspecific scale across habitats within the genus *Helianthus* (Mason and Donovan, 2015). The LHS scheme also appears to be a useful paradigm for understanding divergence, given the high importance of plant height and seed traits alongside leaf economics traits (Westoby, 1998). The smaller stature of desert genotypes is consistent with earlier reproduction during a short, water-limited growing season, with faster leaf economics strategy to support rapid growth in a compressed annual lifespan. Strong diversifying selection on phenology has been detected across populations of the related desert-dwelling *H. anomalus* (Brouillette et al., 2014) as well as across a north-south cline in eastern Great Plains populations of *H. maximiliani* (Kawakami et al., 2011) and *H. annuus* (Blackman et al., 2011), along with accompanying diversifying selection on plant stature, growth rate, or leaf ecophysiology in these species. This evidence from Fst-Qst studies suggests that parallel selective forces may be at work within *H. annuus* between ecoregions. The larger seeds observed in desert genotypes also fits with the LHS paradigm of higher maternal resource provisioning of offspring in habitats with more intense environmental stressors, a strategy to increase early-stage seedling survival (Westoby et al., 1996). Overall, the phenotypic traits identified as the most predictive of ecoregion origin in this dataset are primarily those of the LES and LHS paradigms. This finding validates the importance of these classes of traits to microevolutionary divergence in plant form and function within *H. annuus*, despite the fact that these paradigms were originally developed to describe interspecific variation at community and global scales and their relevance has been repeatedly questioned at smaller scales (Edwards et al., 2014; Niinemets, 2015; Anderegg et al., 2018). Our results also highlight the utility of trait-first ML approaches to test the relative importance of general functional trait paradigms to specific taxa.

The observed importance of floral traits in predicting common sunflower ecoregion origin is less anchored in classical plant functional trait paradigms (but see Roddy et al., 2020 for developing theory). Despite this, floral traits are of course long-understood within plant evolutionary biology to be of critical importance for local adaptation as well as shaped by sexual selection (Darwin, 1862; Stebbins 1970). At the intraspecific scale, variation in floral traits among populations has been demonstrated to be influenced by variation in local pollinator composition (Herrera 2005; Gomez et al., 2014a; Johnson, 2010), and the selection imposed by even generalist pollinators can be very strong for self-incompatible species like *H. annuus* (Galen, 1999). Selection from the abiotic environment is also known to play a role in the evolution of floral traits, as flowers require substantial nutrient investment and are a source of major water loss for plants (Galen, 1999; Teixido et al., 2014). Here we observe that floral phenotypes that most predict ecoregion origin include multiple traits related to the size of the composite head, including the sizes of discs, ligules, and phyllaries, with longer average length of phyllaries being predictive of desert origin. Across the genus *Helianthus*, the evolution of larger heads has been demonstrated to occur repeatedly under lineage diversification into more arid environments (Mason et al., 2017), and assessment of intraspecific divergence in *Helianthus maximiliani* indicates that disc and ligule size are under diversifying selection across climatic gradients with larger structures evolving in drier environments (Kawakami et al. 2011). These similarities across species and taxonomic scales suggest generalities in the evolution of floral traits, and highlight that machine learning approaches like those deployed here can efficiently detect otherwise subtle patterns of intraspecific trait divergence and could be leveraged to advance synthesis and theory development in plant functional trait evolution.

Meta-analysis has estimated that around one-third of all variation in plant functional traits among communities is attributable to intraspecific trait variation, with the remainder attributable to species turnover (Siefert et al., 2015). This means that variation across the landscape within species due to either local adaptation or phenotypic plasticity is substantial, and there is an emerging consensus that axes of differentiation as well as trait-trait relationships at the intraspecific scale often do not reflect global interspecific patterns (Edwards et al., 2014; Niinemets, 2015; Mason and Donovan, 2015; Anderegg et al., 2018). Machine learning methods like the classification approaches applied here have the exciting potential to identify whether there are indeed general patterns of intraspecific trait divergence that occur in parallel in widespread species that span landscape-level environmental gradients. For example, imagine the exact analysis performed here applied to the thousands of plant species that occupy both the Great Plains and North American Deserts, or indeed any contrasting set of ecoregions. The decades-long project of herbarium digitization holds the potential for generating massive plant trait datasets with a magnitude of intraspecific scope that has been lacking from previous global trait datasets (Heberling, 2022), and the deployment of high-throughput phenotyping and remote sensing in a range of settings promises to generate equally impressive ‘phenomic’ datasets (Arend et al., 2022; Cavender-Bares et al., 2022; Gill et al., 2022; Tao et al., 2022). Given the coming tide of truly “big data” plant trait datasets, there is a growing need for creative analytical approaches to synthesizing trait data in ways that will advance our understanding of plant functional trait evolution. Previous generations of scientists rarely had access to datasets of this scope, and we need to identify efficient methods to explicitly test the transferability of traditional paradigms across scales, as well as approaches to generate new insights. In particular, there is major potential to use granular intraspecific-scale datasets to get far closer to the scale at which phenotypic microevolution actually occurs, assessing the multivariate trait divergences among populations that lead to incipient speciation and the eventual subsequent interspecific differences reflected in modern global plant functional trait databases (e.g., Kattge et al., 2020).

## Acknowledgements

The authors would like to acknowledge the massive amount of effort that produced the public HeliantHome database on which this work relies. The authors would like to thank Emily E. Karwacki for the invaluable input with the figures. The authors would also like to thank Rebekah Davis and Charles Pitsenberger for their valuable feedback. Financial support was provided by the Foundation for Food and Agriculture Research under award FF-NIA20— 0000000023. The content of this publication is solely the responsibility of the authors and does not necessarily represent the official views of the Foundation for Food and Agriculture Research.

## Author Contributions

SM and CMM designed the study. SM wrote all code and conducted all analyses. SM created figures and wrote the manuscript with input from CMM.

## Data Availability

All data used in this study, is available from the HeliantHome database (http://www.helianthome.org/). All code used in this analysis as well as the data can be found here (GitHub repository: https://github.com/SamMajumder/Applying-XAI-approaches-to-ecology/tree/master)

---

**Figure S1.** Percentage of missing value for each functional trait in our dataset.

**Figure S2.** Visualizations of the predictions made by the Random Forest (RF) classifier, where points represent populations of *H. annuus*. Each facet represents populations where a given proportion of genotypes was correctly predicted, arranged in descending order – all genotypes.

## Appendices

**Table S1.** Functional trait data from the populations used in this study along with their geographical coordinates and ecoregion membership.

**Table S2.** Training dataset, randomly divided to contain 70% of the individuals (genotypes) and missing data was imputed using proximity to random forest method.

**Table S3.** Testing dataset, containing the remaining 30% of the individuals (genotypes) and missing data was imputed using proximity to random forest

**Table S4.** The optimal subset of ecologically relevant traits which were identified by recursive feature elimination and are ranked based on their importance on a relative scale.

**Table S5.** The traits that were not part of the optimal subset of relevant traits i.e., traits discarded during recursive feature elimination.

**Table S6.** The most ecoregion-divergent traits identified by Boruta and are ranked based on their importance on a relative scale.

**Table S7.** Random forest predicted versus actual ecoregions for each individual genotype in the testing dataset as well as the populations to which they belong to.

**Table S8.** Gradient boosting machine predicted versus actual ecoregions for each individual genotype in the testing dataset as well as the populations to which they belong to. correctly predicted, four out of five, three out of four, two out of three, one out of two, one out of four, and no genotypes correctly predicted by RF.

